# A complete reference genome assembly and annotation of the Black Redstart Phoenicurus ochruros

**DOI:** 10.1101/2025.05.21.655187

**Authors:** Prashant Ghimire, Nan Wang, Sangeet Lamichhaney

**Affiliations:** Department of Biological Sciences, Kent State University, Kent, Ohio, USA; School of Ecology and Nature Conservation, Beijing Forestry University, Beijing, China; School of Biomedical Sciences, Kent State University, Kent, OH 44240, USA

## Abstract

The Black Redstart (*Phoenicurus ochruros*) is one of the most widely distributed species, occupying diverse habitats and exhibiting remarkable altitudinal migration, making it suitable model for studying altitudinal migration and high-altitude adaptation. In this study, we present the first reference genome of *Phoenicurus ochruros*, generated using PacBio HiFi long-read sequencing. The nuclear genome is 1.37 Gb in length, assembled into 296 scaffolds with a scaffold N_50_ of 29.9 Mb. Multiple complementary genome validation approaches - including BUSCO analysis, mapping of raw PacBio HiFi reads, and comparative genomics analysis - confirmed the high contiguity and gene-space completeness of the genome. The Black Redstart genome contains one of the highest proportions of transposable elements among passerines (30.58%), second only to Bell’s Sparrow (*Artemisiospiza belli)*, 31.2%. We further integrated RNA-seq data, protein homology evidence, and *ab initio* gene prediction to build high-quality genome annotation of 18,609 genes in the *P. ochruros* genome. This genomic resource provides invaluable resource for evolutionary genomics studies of passerines and explore genetic basis of high-altitude adaptation.

## Background & Summary

Altitudinal migrant birds are species that move up and down along the elevation gradients - usually within the same geographic region - in response to seasonal changes in climate, food availability, or breeding conditions^1^. Unlike long-distance migrants that travel between continents or across hemispheres, altitudinal migrants move vertically between lowland and highland areas, often over relatively short horizontal distances. The majority of birds residing in mountain ecosystems undergo seasonal altitudinal migration, experiencing rapid changes in climate, habitat, and oxygen availability, which can shape their patterns of local adaptation and drive their population divergence^2^. This form of migration allows them to exploit seasonal resources while avoiding harsh environmental conditions. Notably, more than ∼65% of bird species in the Himalayas are altitudinal migrants, and they offer a unique model system for studying patterns of species divergence, mechanisms of rapid physiological acclimation, and long-term genetic adaptation to both short- and long-term environmental changes^3^.

One well-known example of altitudinal migrants are the Redstarts (Family: Muscicapidae, genus: *Phoenicurus*), which exhibit seasonal elevational shifts in the Himalayas, moving to higher elevations in summer for breeding and descending to lower elevations in winter (**Fig. 1a**).

**Figure 1:**
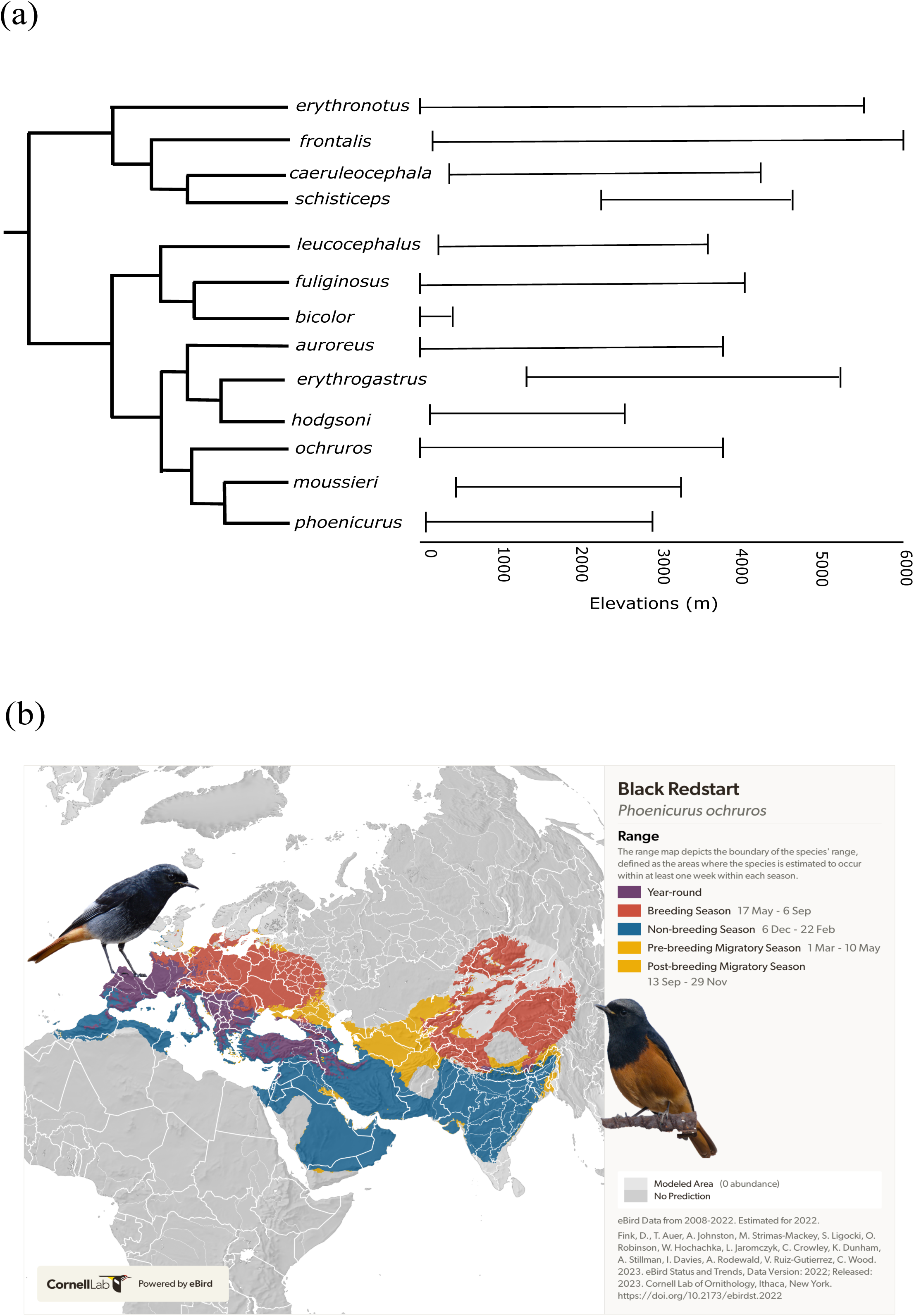
Redstarts (Genus: *Phoenicurus*) distribution across the altitudinal gradient. **(a)** Phylogeny^38^ and altitudinal range^39^ show significant variation in the altitudinal distribution of redstarts **(b).** The seasonal distribution range of the Black Redstart *(Phoenicurus ochruros)* spans across Asia, Europe, and Africa^6^. The Eastern Black population (right; © Satdeep Gill, Wikimedia Commons) and Western Black population (left; © Rabiruivo comum, Wikimedia Commons) exhibit contrasting plumage patterns on their bellies, highlighting the geographical variation within the species.

Redstarts are a group of 14 species of small Old-World flycatchers, native to Asia, Europe and Africa^4^, known for their striking plumage and vocalization diversity^5^. Among them, the Black Redstart (*Phoenicurus ochruros*), distinguished by its dark plumage and orange lower belly, is one of the most widely distributed Redstarts, ranging from western Europe and North Africa to Asia^5^ (**Fig. 1b**). It thrives in a variety of habitats, including shrublands, grasslands, rocky slopes up to the snow line, and urban areas near human settlements^5,6^. It typically breeds above the tree line (>4000 m) and migrates down to lower elevation during the non-breeding winter season (0-2500 m)^5^. Black Redstarts exhibit substantial morphological variation across its range, with eight subspecies currently recognized based on the variation in phenotypic traits^6^ (**Fig. 1b**). Broadly, individuals found in Europe and northern Africa with grey and white bellies are referred to as Western Black Redstarts, while those with orange bellies found across Asia are known as Eastern Black Redstarts^6^. Although these birds show clear morphological differences, they are classified as a single species due to the presence of intermediate populations in the Middle East that exhibit transitional plumage, vocalizations, and mitochondrial DNA^7,8^. However, this taxonomic treatment remains unresolved, largely due to the lack of comprehensive genomic studies on the species^6^.

Genomic studies are essential in Black Redstarts because current subspecies designations are based primarily on phenotypic traits - such as plumage coloration and vocalizations and limited molecular markers - that can be shaped by both genetics and environment. While intermediate populations suggest possible gene flow between Eastern and Western populations, genomic data can characterize the magnitude and direction of this gene flow, helping to determine whether these forms represent (a) distinct evolutionary lineages (potentially cryptic species), (b) recently diverged populations still connected by hybrid zones, or (c) subspecies undergoing incomplete lineage sorting. Genomic analyses can identify patterns of selection, identify regions of the genome associated with phenotypic divergence between populations, and reveal how historical processes like glaciations or range shifts have shaped genetic structure across their distribution range. Most importantly, Black Redstarts offer a promising model system for investigating the genetic basis of altitudinal migration. Their seasonal movements are associated with rapid changes in temperature, oxygen availability, and resource distribution - conditions that are known to impose strong selective pressures on metabolic, cardiovascular, and behavioral traits^9,10^. By studying the genomes of altitudinal migrants like Black redstarts, we can identify genetic variants and regulatory pathways linked to altitude-related physiological adaptations, such as hypoxia tolerance, energy metabolism, and timing of migration.

In this study, we present the first high-quality genome assembly and annotation of the Black Redstart, generated using a combination of long-read PacBio HiFi sequencing and Illumina short-read RNA sequencing. Our efforts resulted in the successful assembly of both the nuclear and mitochondrial genomes, marking the first time a reference genome has been produced for any species in the genus *Phoenicurus*. This reference genome provides a valuable genomic resource for future analyses of population demographic history and biogeographic patterns, and facilitates studies on the genetic basis of high-altitude adaptation. Furthermore, it lays the groundwork for comparative genomics across the Muscicapidae family, advancing our understanding of avian diversification, adaptation, and migration at both macro- and microevolutionary scales.

## Methods

### Sample collection

An adult female Black Redstart *(Phoenicurus ochruros)* was captured using mist nets at an elevation of 3,000 meters in the Qinghai-Tibetan Plateau (QTP). Blood sample was collected via brachial venipuncture and immediately stored in RNAlater until further processing for DNA extraction. Following blood sampling, the bird was humanely euthanized following institutional animal care guidelines. The individual was then dissected, and tissue samples were collected from the brain, liver, lungs, gonads, spleen, kidney, heart, and pectoral muscle. All tissues were preserved in RNAlater for subsequent RNA extractions. All procedures were conducted under approved animal care protocols and valid collection permits.

### DNA extraction and genome sequencing

Genomic DNA was extracted from tissue samples using the DNeasy Blood & Tissue Kit (Qiagen, Valencia, CA, USA) following the manufacturer’s protocol. DNA quality and quantity were assessed using spectrophotometry and gel electrophoresis to ensure high molecular weight suitable for long-read sequencing. Long-read whole-genome sequencing was performed on the PacBio Revio platform, which generated highly accurate HiFi (high-fidelity) reads, enabling robust assembly of complex genomic regions. We generated a total of ∼93 Gb of raw PacBio HiFi long reads with an average read length of 17.39 Kb and read N50 of 17.41 kb (**Fig. 2a**).

**Figure 2:**
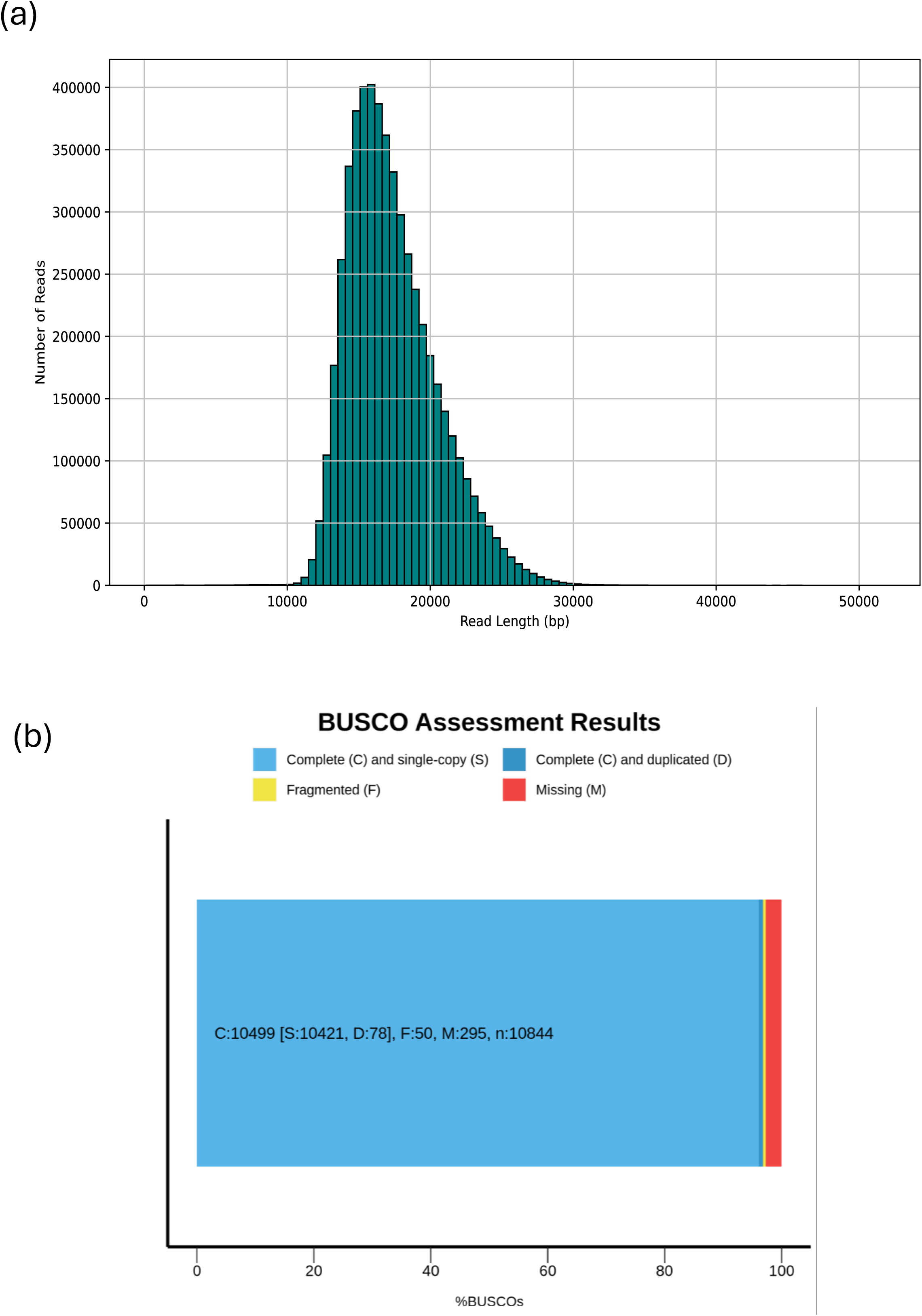
Statistics on raw PacBio HiFi long reads and de novo genome assembly. **(a)** Length distribution of PacBio HiFi raw reads **(b)** BUSCO (Benchmarking Universal Single-Copy Orthologs) assessment of genome completeness for *Phoenicurus ochruros*. The results indicate that 96.8% of BUSCO genes were identified as complete, including 96.1% complete and single-copy and 0.7% duplicated BUSCOs. Only 2.2% were fragmented and 1.0% were missing, demonstrating high completeness and gene-space coverage of the assembly.

### Genome assembly and quality assessment

We assembled ∼93 Gb of raw PacBio HiFi long reads using HiFiasm (v.0.24.0) - a fast and accurate assembler optimized for PacBio HiFi data^11^. We used the default parameters in HiFiasm with built-in duplication purging. The resulting *de-novo* genome assembly of *P. ochruros* was 1.37 Gb in total length, consisting of 296 scaffolds, with the Scaffold N_50_ of 29.9 Mb and a longest scaffold of length of 113.3 Mb (**Table 1**). These results indicated a high level of genome contiguity. Furthermore, the completeness of genome assembly was evaluated using BUSCO (v.5.7.1)^12^ against the passeriformes_odb10 database.

The results revealed 96.80% of BUSCOs were successfully detected, including 96.10% complete and single copy, and 0.7 % of duplicated BUSCOs (**Table 1**, **Fig. 2b**, **Fig. 3a**), indicating the high level of genome completeness. In addition, we mapped the raw HiFi reads back to the assembled genome using Minimap2 (v.2.21)^13^, and found that 100% of the reads successfully mapped, further supporting the high completeness and accuracy of the genome assembly.

**Figure 3:**
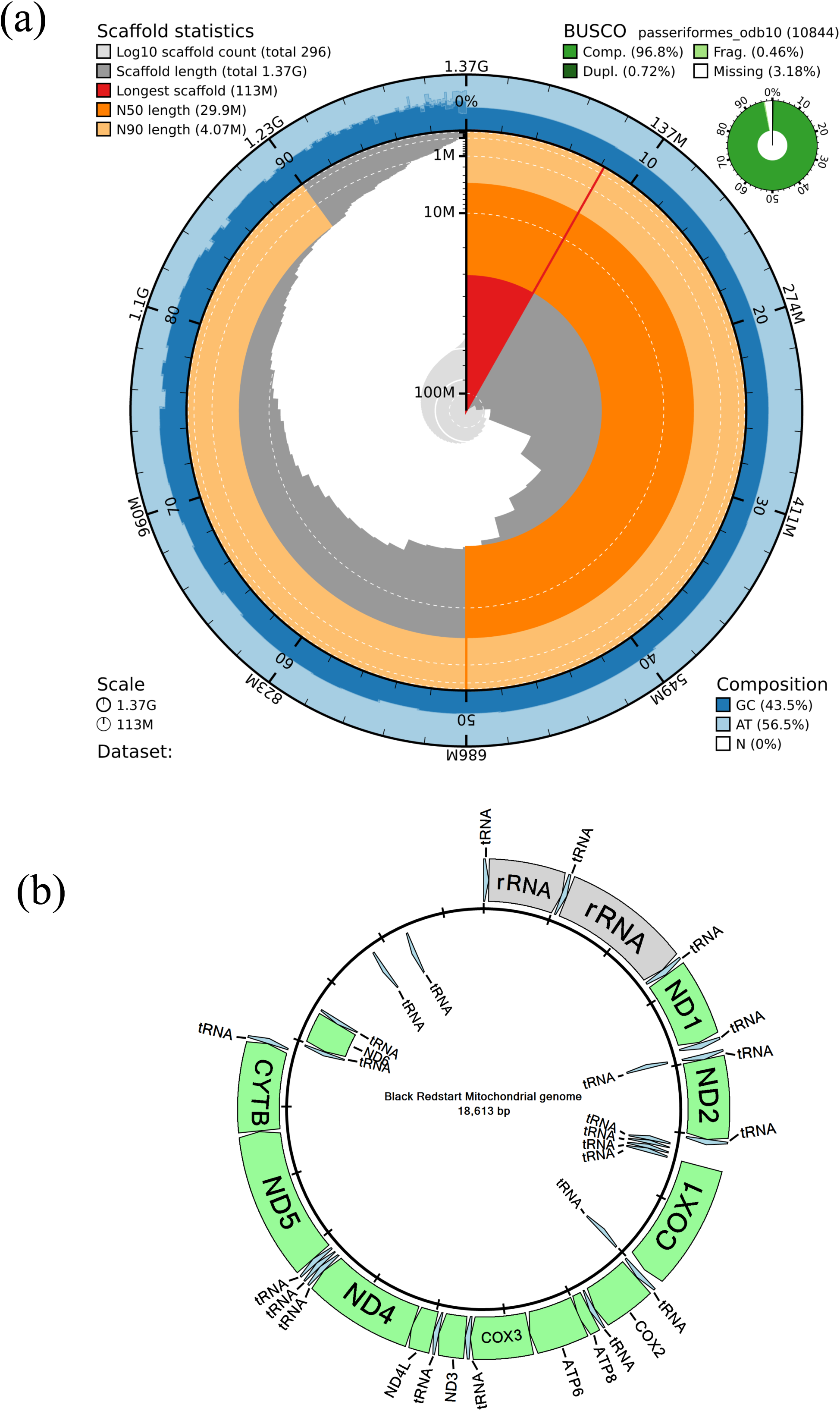
Summary of Genome assembly statistics. **(a)** Statistics of the nuclear Genome. The distribution of sequence lengths is shown in dark grey with the plot radius scaled to the longest sequence present in the assembly (shown in red). Orange and pale-orange arcs show the N50 and N90 sequence lengths, respectively. The pale grey spiral shows the cumulative sequence count on a log scale with white scale lines showing successive orders of magnitude. The blue and pale-blue area around the outside of the plot shows the distribution of GC, AT, and N percentages in the same bins as the inner plot. A summary of complete, fragmented, duplicated, and missing BUSCO genes in the passeriformes_odb10 set is shown in the top right **(b)** Details of the mitochondrial genome of the Black redstart showing position of genes and tRNA.

### Mitochondrial genome assembly and annotation

We assembled the mitochondrial genome using the MitoHiFi v3.0.0 pipeline^14^, which is optimized for extracting and assembling mitochondrial sequences from PacBio HiFi reads. We applied the parameter setting of “-p 90”, which retained contigs with 90% or more of their length matching a reference mitochondrial sequence in BLAST searches. As a reference, we used the published mitochondrial genome of the closely related Daurian Redstart *(Phoenicurus auroreus)* (GenBank accession: MT366880.1^15^). This approach enabled the successful assembly of a complete mitochondrial genome for *P. ochruros*. The final draft mitochondrial genome assembly of the *P. ochruros* was 18,613 bp in length and includes 13 protein-coding genes, 24 tRNA genes, and two ribosomal RNA genes, consistent with the typical structure of vertebrate mitogenomes (**Fig. 3b**).

### Repeat annotation and masking

Before predicting protein-coding genes in the genome, repeat annotation was conducted using a combination of *de novo* and homology-based approaches to identify and mask repetitive elements in the genome. We first used RepeatModeler (v.2.0.5)^16^ to build a *de novo* custom-built species-specific repeat library generated for *P. ochruros* to increase the accuracy of detection and annotation of transposable elements. This custom repeat library was then merged with the reference repeat library downloaded from the Repbase library (release 20190301)^17^ for identification and masking of transposable elements in the *P. ochruros* genome using RepeatMasker (v.4.1.6)^18^.

We identified 30.58% of the Black Redstart genome as transposable elements. These included 0.13% Short Interspersed Nuclear Elements (SINEs), 5.44% Long Interspersed Nuclear Elements (LINEs), 7.75% Long Terminal Repeat (LTR) elements, 0.17% DNA transposons, and 0.10% rolling-circle elements (**Table 2**). Additionally, 4.37% of the genome consisted of unclassified repeats, and a notable 10.77% was composed of satellite DNA. Such a high proportion of transposable elements is uncommon among birds^19^. Notably, only a few passerines - such as Bell’s Sparrow *(Artemisiospiza belli)*, which exhibits a repeat content of 31.2%^20^ display similarly elevated levels of repetitive elements. The genome size of the Black Redstart is comparable to that of Bell’s Sparrow (**Fig. 4a**, **Table 2**), and since repeat content is generally correlated with genome size^19^, the higher proportion of repetitive elements in the Black Redstart genome is expected. In contrast, a close relative within the Muscicapidae family, the Spotted Flycatcher *(Muscicapa striata)*, possesses a smaller genome size of 1.08 Gb and a significantly lower transposable element content of only 13.97%^21^ (**Fig. 4a**, **Table 2**). Compared to Bell’s Sparrow *(Artemisiospiza belli)*, which has 14.5% unclassified repeats, our repeat annotation for *Phoenicurus ochruros* is more complete, with only 4.37% of repeats remaining unclassified. The high repeat content observed in *P. ochruros* suggests that its genome may serve as a valuable resource for investigating the evolutionary dynamics and regulatory roles of transposable elements in songbirds.

**Figure 4:**
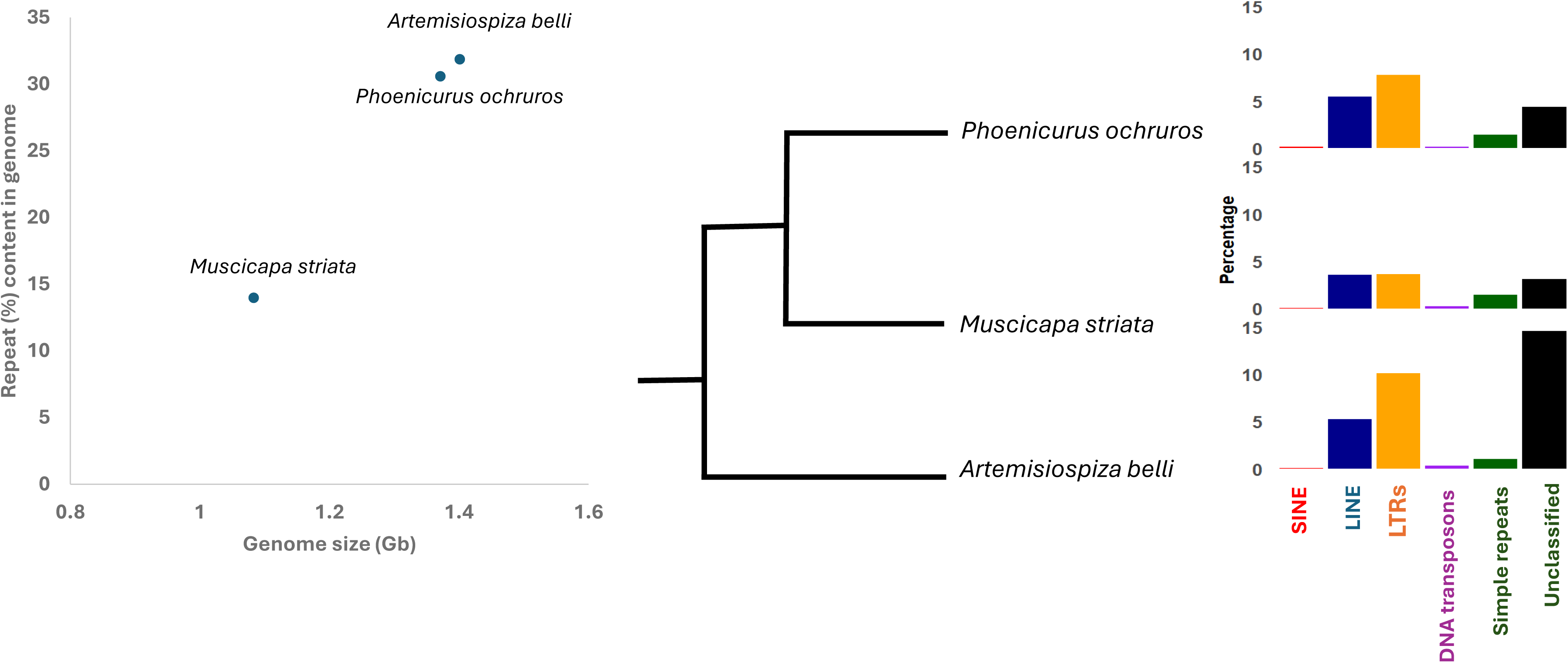
Landscape of Transposable Elements (TE) in *Phoenicurus ochruros, Artemisiospiza belli,* and *Muscicapa striata* (a) Correlation of TE content and genome size **(b)** Comparison of various categories of TE across three species.

### RNA extraction and transcriptome sequencing

Total RNA was extracted from eight tissue types - brain, liver, lungs, gonads, spleen, kidney, heart, and pectoral muscle - using the RNeasy Mini Kit (Qiagen, Valencia, CA, USA), following the manufacturer’s protocol. RNA quality and integrity were assessed using a Bioanalyzer and spectrophotometry to ensure suitability for downstream sequencing. Each tissue sample was sequenced on the Illumina NovaSeq platform with 150bp paired-end reads, generating approximately 45 million paired-end RNA-seq reads (total ∼ 6.5 Gb of data) per sample (**Table 3**).

### Genome annotation

We performed gene prediction and annotation using BRAKER3, an automated pipeline that integrates RNA-seq data, protein homology, and *ab initio* gene prediction to generate high-confidence gene models^22^. BRAKER3 runs the GeneMark-ETP pipeline^23^, which executes a series of steps designed to accurately predict protein-coding genes. First, RNA-seq reads from eight tissues were aligned to the genome using HISAT2^24^. These alignments were used by StringTie2^25^ to assemble transcript sequences, which were then analyzed using GeneMarkS-T^26^ to identify putative protein-coding genes. The resulting protein sequences were searched against a reference vertebrate protein OrthoDB11^27^ database to determine similarity scores and filter out high-confidence gene predictions. The parameters of GeneMark.hmm were then trained on this high-confidence gene set, which was used to predict genes in the intermediate, less well-supported genomic regions. These intermediate predictions were matched against homologous proteins in the OrthoDB database, and the alignments were mapped back to the genome using ProtHint^28^ to generate hints (e.g., intron-exon boundaries, start/stop codons). Hints were classified into three groups: (a) Transcript and protein similarity support, (b) Transcript and *ab initio* support, and (c) Protein similarity support only. All three sets were used to expand the initial high-confidence gene set into a genome-wide gene annotation. Next, AUGUSTUS (v3.5.0)^29^ was trained on the high-confidence gene models to generate a second set of gene predictions, integrating the RNA-seq and protein-derived hints. To further improve the accuracy of gene prediction, the *Passeriformes_odb10* lineage dataset from BUSCO^30^ was used to optimize AUGUSTUS parameters by training on evolutionarily conserved genes. This ensured that gene models were informed by conserved genomic features specific to passerine birds, resulting in enhanced prediction quality. Finally, TSEBRA, a tool integrated within BRAKER3, was used to combine the gene predictions generated by GeneMark-ETP and AUGUSTUS. TSEBRA ensured the inclusion of high-confidence genes in the final annotation set and selects optimal gene models based on a scoring system that prioritizes evidence-supported predictions. This comprehensive approach resulted in a high-quality, evidence-supported annotation of 18,609 genes in the *P. ochruros* genome.

Functional annotation of protein-coding genes was carried out using BLASTp (v2.13.0+)^31^ against the Swiss-Prot database (UniProt release 2024_07)^32^, applying an e-value threshold of 1 × 10. Protein domains were identified by querying the Conserved Domain Database (CDD) and Pfam^33^ using InterProScan (v.5.67-99.0)^34^. A total of 18,609 gene models were identified, with an average gene length of 23,222 bp, approximately 20 exons per gene and with average exon length of 163 bp (**Fig. 5**). Gene Ontology (GO) terms and Kyoto Encyclopedia of Genes and Genomes (KEGG) annotation for the genome were obtained using eggNOG-mapper (v.2.0)^35^ pipeline. We further evaluated the completeness of annotated protein sequences using BUSCO (v. 5.7.1)^12^ against the *passeriformes_odb10* database. 97.8% of BUSCOs were successfully detected, including 96.6% complete and single copy, and 1.2% of duplicated BUSCOs. These results were consistent with the BUSCO assessment results based on the genome assembly (**Table 1**, **Fig. 2b**).

**Figure 5:**
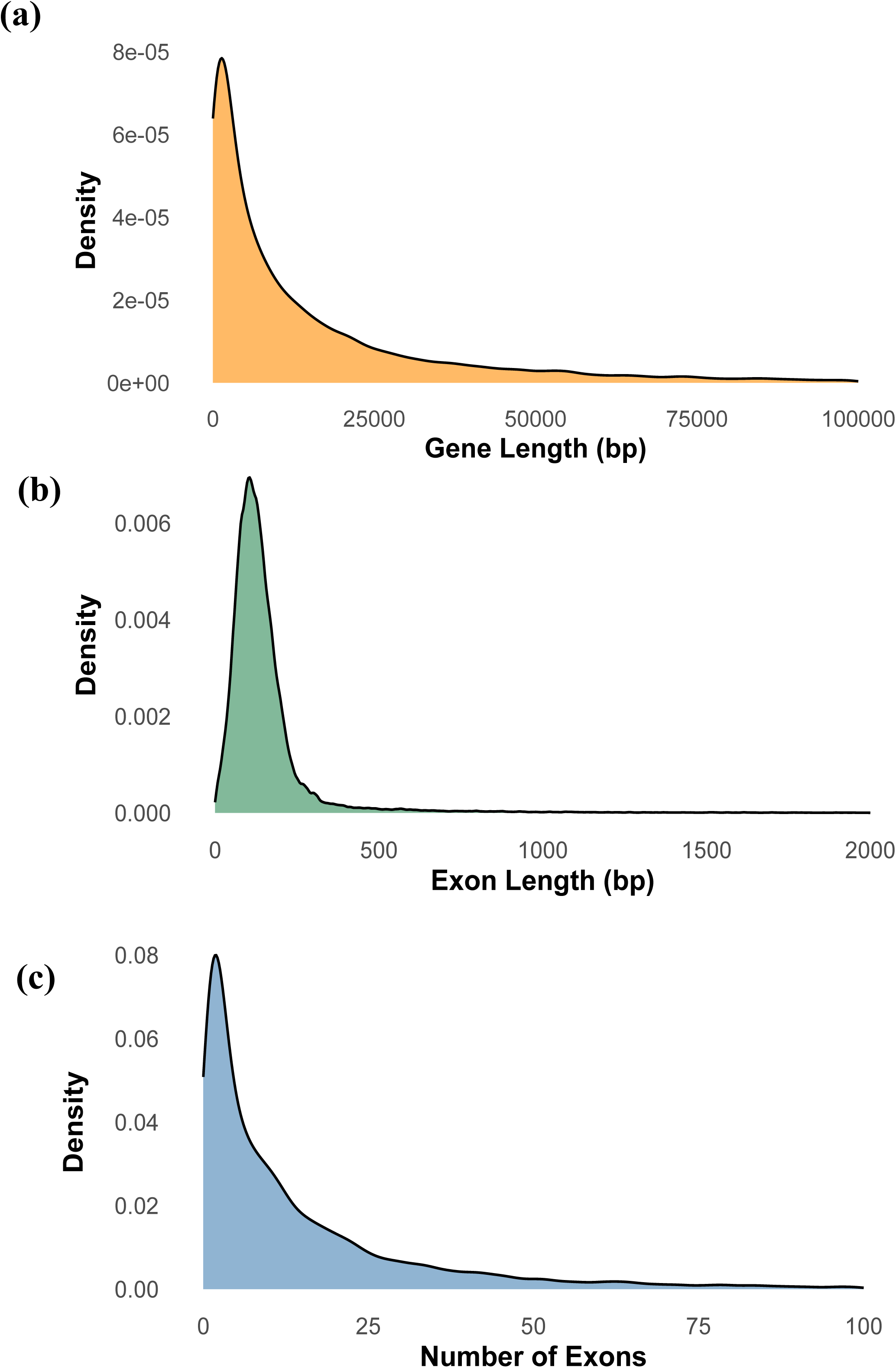
Gene structure statistics of *Phoenicurus ochruros*. **(a)** The distribution of gene length **(b)** The distribution of exon length, and **(c)** The distribution of the number of exons per gene.

### Data Records

The PacBio HiFi long reads and the RNAseq data have been deposited in the Sequence Read Archive (SRA) database with accession number SRP584347 under BioProject accession PRJNA121710 (https://www.ncbi.nlm.nih.gov/bioproject/PRJNA1217107). The genome assembly has been submitted to NCBI GenBank under the accession JBLKNL000000000 (https://www.ncbi.nlm.nih.gov/nuccore/JBLKNL000000000). The genome annotation files are available in Figshare (https://doi.org/10.6084/m9.figshare.28912433). The mitochondrial genome and its annotation have been submitted to NCBI GenBank under the accession ID PV606442 (https://www.ncbi.nlm.nih.gov/nucleotide/PV606442).

### Technical Validation

DNA and RNA purity were assessed using a NanoDrop 2000 Spectrophotometer (Thermo Fisher Scientific, Waltham, MA, USA). DNA and RNA concentration were quantified with a Qubit 2.0 Fluorometer (Thermo Fisher Scientific), and DNA/RNA integrity was evaluated using an Agilent 4200 Bioanalyzer (Agilent Technologies, Santa Clara, CA, USA). Only high-quality DNA and RNA samples were selected for genome and transcriptome sequencing. To validate the quality of the assembled genome, we employed multiple complementary approaches. First, we evaluated genome contiguity and gene-space completeness using standard assembly statistics and BUSCO^12^. The assembly achieved a scaffold N_50_ of ∼30 Mb and a BUSCO completeness score of 97.30% against the passeriformes_odb10 database, indicating a high contiguity and gene-space completeness.

Repeat-rich regions are often difficult to assemble accurately due to their repetitive nature, making them a common source of assembly errors or gaps. By evaluating the abundance and composition of transposable elements among representative species with published genomes, we can validate the quality of a *de novo* genome assembly. We compared the repeat landscape of *P. ochruros* with that of several other vertebrate species (**Table 2**); Blind cave fish *(Astyanax mexicanus)* (GenBank accession: GCA_023375975.1), Common frog (*Rana temporaria)* (GenBank accession: GCA_905171775.1), Chicken (*Gallus gallus)* (GenBank accession: GCA_016699485.1), Bell’s sparrow (*Artemisiospiza belli)* (GenBank accession: PRJNA720569), Nelson’s sparrow (*Ammospiza nelsoni)*^36^, White-throated sparrow *(Zonotrichia albicollis)* (GenBank accession: PRJNA197293), Spotted flycatcher (*Muscicapa striata)*^21^, Obi paradise-crow (*Lycocorax pyrrhopterus obiensis)*^37^, and White wagtail (*Motacilla alba)* (GenBank accession: GCA_015832195.1). Our results indicated that the repeat content in *P. ochruros* is consistent with known evolutionary trends and genomic patterns observed in other passerines. We found a strong positive correlation between genome size and repeat content (*r* = 0.87), indicating that the assembly has captured repetitive elements with high accuracy. This comparative approach not only aids in identifying potential assembly artifacts, such as collapsed or missing repeat regions, but also supports the completeness of repeat annotation. Together, these findings enhance confidence in the overall quality and biological validity of the *Phoenicurus ochruros* genome assembly.

## Code availability

All custom scripts and codes used in this study are available on GitHub at - https://github.com/Prashantevo/Black_redstart_genome_assembly.

## Acknowledgements

The fieldwork was supported by the National Survey on Terrestrial Wildlife Resources in China to NW. PG was supported by a graduate assistantship from Kent State University. Genome sequencing was supported from Kent State University to SL.

## Authors contributions

SL designed the study, NW collected the samples, PG conducted the bioinformatics analysis under SL supervision. PG and SL drafted the manuscript. All authors read and approved the final version of the manuscript.

