## Supplementary material for "A complete reference genome assembly and annotation of the Black Redstart Phoenicurus ochruros": Table

**Table 1: Genome assembly statistics**

| Total genome size | 1,371,345,233 bp |
| --- | --- |
| Minimum length of scaffold | 17,018 bp |
| Maximum length of scaffold | 113,313,014 bp |
| Number of Scaffolds | 296 |
| Average length of scaffold | 4,632,923 bp |
| Scaffold N50 | 29,902,800 bp |
| BUSCO Assessment Statistics (using *Passeriformes_odb10* dataset) |  |
| C (Complete) | 96.80% |
| S (Complete and Single-copy) | 96.10% |
| D (Complete and Duplicated) | 0.70% |
| F (Fragmented) | 0.50% |
| M (Missing) | 2.70% |
| Total BUSCO Groups Searched | 10,844 |
| Complete BUSCOs (C) | 10,499 |
| Complete and Single-copy BUSCOs (S) | 10,421 |
| Complete and Duplicated BUSCOs (D) | 78 |
| Fragmented BUSCOs (F) | 50 |
| Missing BUSCOs (M) | 295 |

**Table 2: Repeats summary for Black redstart and other selected species**

| Feature | Black Redstart | Bell's Sparrow | Chicken | Mexican Tetra Fish | Nelson's Sparrow | Paradise Crow | Spotted Flycatcher | White Wagtail | White-throated Sparrow |
| --- | --- | --- | --- | --- | --- | --- | --- | --- | --- |
| Genome size | 1371345233 | 1401818823 | 1.05E+09 | 1373169186 | 1185463352 | 1107771238 | 1083165776 | 1072670728 | 1052600561 |
| Log (genome_size) | 9.137146801 | 9.146691887 | 9.022565 | 9.137724049 | 9.073888133 | 9.044450085 | 9.034694929 | 9.030466429 | 9.022263597 |
| Bases Masked (%) | 30.58 | 31.86 | 14.59 | 53.85 | 20.11 | 12.29 | 13.97 | 12.34 | 8.89 |
| SINE (%) | 0.13 | 0.07 | 0.09 | 0.08 | 0.08 | 0.09 | 0.08 | 0.08 | 0.08 |
| Retroelements (%) | 13.32 | 15.37 | 8.5 | 5.75 | 10.06 | 7.05 | 7.29 | 5.46 | 5.14 |
| LINEs (%) | 5.44 | 5.22 | 7.4 | 3.32 | 6.17 | 4.25 | 3.58 | 3.78 | 3.02 |
| LTR Elements (%) | 7.75 | 10.08 | 1.02 | 2.35 | 3.81 | 2.71 | 3.62 | 1.6 | 2.05 |
| DNA Transposons (%) | 0.17 | 0.36 | 0.98 | 20.97 | 0.26 | 0.31 | 0.24 | 0.28 | 0.27 |
| Rolling Circles (%) | 0.1 | 0.04 | 0 | 3.31 | 0.03 | 0.01 | 0.13 | 0.02 | 0.02 |
| Unclassified (%) | 4.37 | 14.55 | 3.38 | 19.91 | 8.07 | 3.16 | 3.14 | 5.1 | 5.2 |
| Total Interspersed Repeats (%) | 17.86 | 30.28 | 12.86 | 46.63 | 18.39 | 10.53 | 10.66 | 10.83 | 9.23 |
| Small RNA (%) | 0.22 | 0.03 | 0.06 | 0.36 | 0.07 | 0.28 | 0.09 | 0.02 | 0.01 |
| Satellites (%) | 10.77 | 0.27 | 0.04 | 0.77 | 0.2 | 0.49 | 1.34 | 0.22 | 0.18 |
| Simple Repeats (%) | 1.41 | 1.06 | 1.34 | 2.46 | 1.15 | 0.79 | 1.46 | 1.02 | 0.95 |
| Low Complexity (%) | 0.23 | 0.21 | 0.3 | 0.33 | 0.29 | 0.21 | 0.3 | 0.24 | 0.22 |

**Table 3: Summary of RNAseq data generated from eight tissues**

| Sample | Tissue | Reads (bp) | Reads (Gb) |
| --- | --- | --- | --- |
| B-7-FE | Lungs | 44,465,278 | 6.669792 |
| B-7-g | Liver | 44,008,762 | 6.601314 |
| B-7-L | Gonads | 48,437,250 | 7.265588 |
| B-7-N | Brain | 47,297,884 | 7.094683 |
| B-7-P | Spleen | 45,808,536 | 6.87128 |
| B-7-SH | Kidney | 41,073,742 | 6.161061 |
| B-7-X | Heart | 44,556,880 | 6.683532 |
| B-7-XJ | Pectoral muscle | 54,275,560 | 8.141334 |
